# Intra-host Variation and Evolutionary Dynamics of SARS-CoV-2 Population in COVID-19 Patients

**DOI:** 10.1101/2020.05.20.103549

**Authors:** Yanqun Wang, Daxi Wang, Lu Zhang, Wanying Sun, Zhaoyong Zhang, Weijun Chen, Airu Zhu, Yongbo Huang, Fei Xiao, Jinxiu Yao, Mian Gan, Fang Li, Ling luo, Xiaofang Huang, Yanjun Zhang, Sook-san Wong, Xinyi Cheng, Jingkai Ji, Zhihua Ou, Minfeng Xiao, Min Li, Jiandong Li, Peidi Ren, Ziqing Deng, Huanzi Zhong, Huanming Yang, Jian Wang, Xun Xu, Tie Song, Chris Ka Pun Mok, Malik Peiris, Nanshan Zhong, Jingxian Zhao, Yimin Li, Junhua Li, Jincun Zhao

## Abstract

As of middle May 2020, the causative agent of COVID-19, SARS-CoV-2, has infected over 4 million people with more than 300 thousand death as official reports^1,2^. The key to understanding the biology and virus-host interactions of SARS-CoV-2 requires the knowledge of mutation and evolution of this virus at both inter- and intra-host levels. However, despite quite a few polymorphic sites identified among SARS-CoV-2 populations, intra-host variant spectra and their evolutionary dynamics remain mostly unknown. Here, using deep sequencing data, we achieved and characterized consensus genomes and intra-host genomic variants from 32 serial samples collected from eight patients with COVID-19. The 32 consensus genomes revealed the coexistence of different genotypes within the same patient. We further identified 40 intra-host single nucleotide variants (iSNVs). Most (30/40) iSNVs presented in single patient, while ten iSNVs were found in at least two patients or identical to consensus variants. Comparison of allele frequencies of the iSNVs revealed genetic divergence between intra-host populations of the respiratory tract (RT) and gastrointestinal tract (GIT), mostly driven by bottleneck events among intra-host transmissions. Nonetheless, we observed a maintained viral genetic diversity within GIT, showing an increased population with accumulated mutations developed in the tissue-specific environments. The iSNVs identified here not only show spatial divergence of intra-host viral populations, but also provide new insights into the complex virus-host interactions.

---

From January 25 to February 10 in 2020, we collected a total of 62 serial clinical samples from eight hospitalized patients (GZMU cohort) confirmed with SARS-CoV-2 infection using real-time RT-qPCR (**Table S1**). All patients had direct contacts with confirmed cases during the early stage of the outbreak. Most patients, except P15 and P62, had severe symptoms and received mechanical ventilation in ICU, including the patient P01 who passed away eventually. The patient P01 also showed much lower antibody (IgG and IgM) responses (**Table S1**) compared to other patients. We then deep sequenced the 62 clinical samples using metatranscriptomic and/or hybrid capture methods (**Table S1**). The numbers of SARS-CoV-2 reads per million (SARS-CoV-2 RPM) among the metatranscriptomic data correlated well with the corresponding RT-qPCR cycle threshold (Ct), reflecting a robust estimation of viral load (R = 0.71, *P* = 6.7e-11) (**Fig. 1a**). The respiratory tract (RT: Nose, Sputum, Throat) and gastrointestinal tract (GIT: Anus, Feces) samples showed higher SARS-CoV-2 RPMs compared to gastric mucosa and urine samples (**Fig. 1b**). Furthermore, RT and GIT samples from two patients with mild symptoms showed relatively low viral loads among their respective sample types. The data here may reflect an active replication of SARS-CoV-2 in RT and GIT, especially in patients with severe symptoms^3,4^.

**Figure 1.**
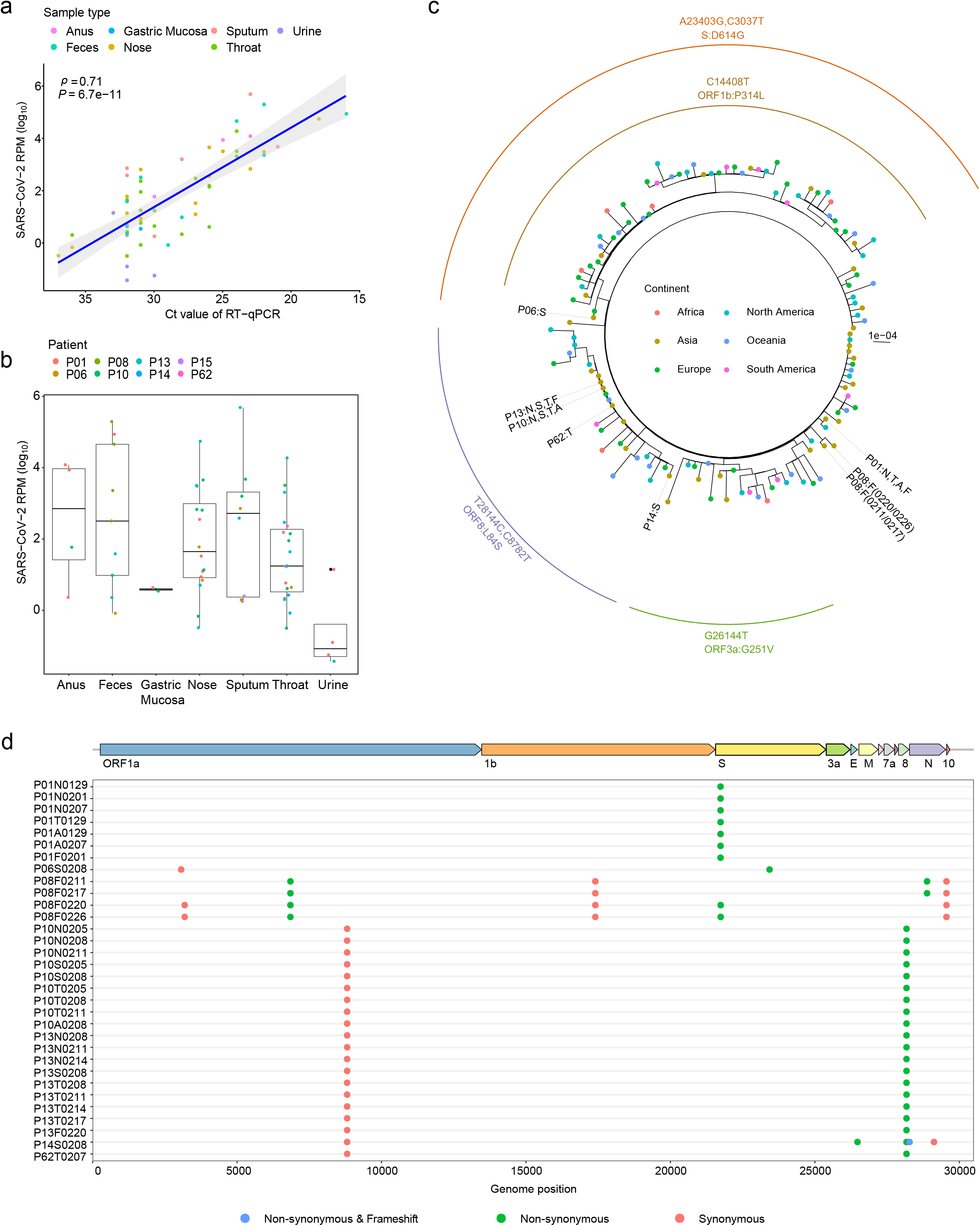
Sequence data from various sample types of patients with COVID-19. **a**, SARS-CoV-2 RPM of meta-transcriptomic data plotted against RT–qPCR cycle threshold (Ct) value for the clinical samples. **b**, Frequency distribution of samples based on SARS-CoV-2 reads per million (SARS-CoV-2 RPM). **c**, Maximum likelihood tree of consensus SARS-CoV-2 genomes using IQ-TREE (1,000 bootstrap replicates). Colors of dotted tips represent geographic locations of samples. Node labels represent bootstrap values for each branch. Nucleotide mutations that defines the branch were labelled outside the tree. **d**, Distribution of consensus variants (in round circles) detected in GZMU cohort across the SARS-CoV-2 genome. Colors represent the biological effect of mutations. Non-synonymous variants are denoted by green, synonymous variants by red, and frameshift by blue. EPI_ISL_402119 was used as the reference sequence.

Here, using metatranscriptomic data, we obtained 32 consensus complete genomes from the clinical samples with at least 60-fold sequence coverage (**Table S1 and Table S2**). Comparing the assemblies to the reference sequence (GISAID accession: EPI_ISL_402119) revealed 14 consensus variants (6 synonymous and 8 non-synonymous) located mostly in ORF1ab, S and N genes (**Table S2**). Most of the consensus variants were also detected among public sequences, including the widespread associated variants (C8782T and T28144C) detected in four patients (P10, P13, P14 and P62). The novel consensus variant causes a frameshift at the end of ORF8 in the patient P14, showing the phenotypic plasticity during the evolution of SARS-CoV-2. Evolutionary relationships showed that the consensus SARS-CoV-2 genomes of the GZMU cohort belonged to distinct clades, including clades defined by T28144C and A23403G, respectively (**Fig. 1c**). Remarkably, we observed distinct SARS-CoV-2 genomes co-existed in the GIT samples of the patient (P08) with three nucleotide differences (**Fig. 1d and Table S2**), suggesting independent replications of different SARS-CoV-2 genotypes within the same host^5^.

Although plenty of polymorphic sites were identified among SARS-CoV-2 populations, intra-host variant spectra of closely related viral genomes are mostly disguised by the consensus sequences. We firstly examined the reproducibility of our experimental procedures for allele frequency identification. Only a minor difference of alternative allele frequencies (AAFs) was observed among biological replicates of two selected samples (**Fig. S1**), showing that the estimated population composition was marginally affected by independent experimental procedures. To control false discovery rate, we applied a stringent approach to detect iSNVs. The iSNVs were identified from the 32 samples using metatranscriptomic data and then verified using hybrid capture data, which are available for most (27/32) samples (**Table S3 and Table S4**). Overall, we observed 1 to 23 iSNVs in six patients with a cut-off of 5% minor allele frequency (**Fig. 2a and Fig. 2b**). When an iSNV was discovered in one patient, we reduced the cut-off to 2% to detect that iSNV from the rest samples of the same patient (see methods). The AAFs of iSNVs detected from the metagenomic data correlated well with those of the hybrid capture data (Spearman’s *ρ* = 0.99, *P* < 2.2e-16; **Fig. S2**). Furthermore, the numbers of the observed iSNVs did not correlate with the sequencing coverage (**Fig. S3**), suggesting that the coverage of metatrancriptomic data was sufficient to estimate intra-host variation in most samples.

**Figure 2.**
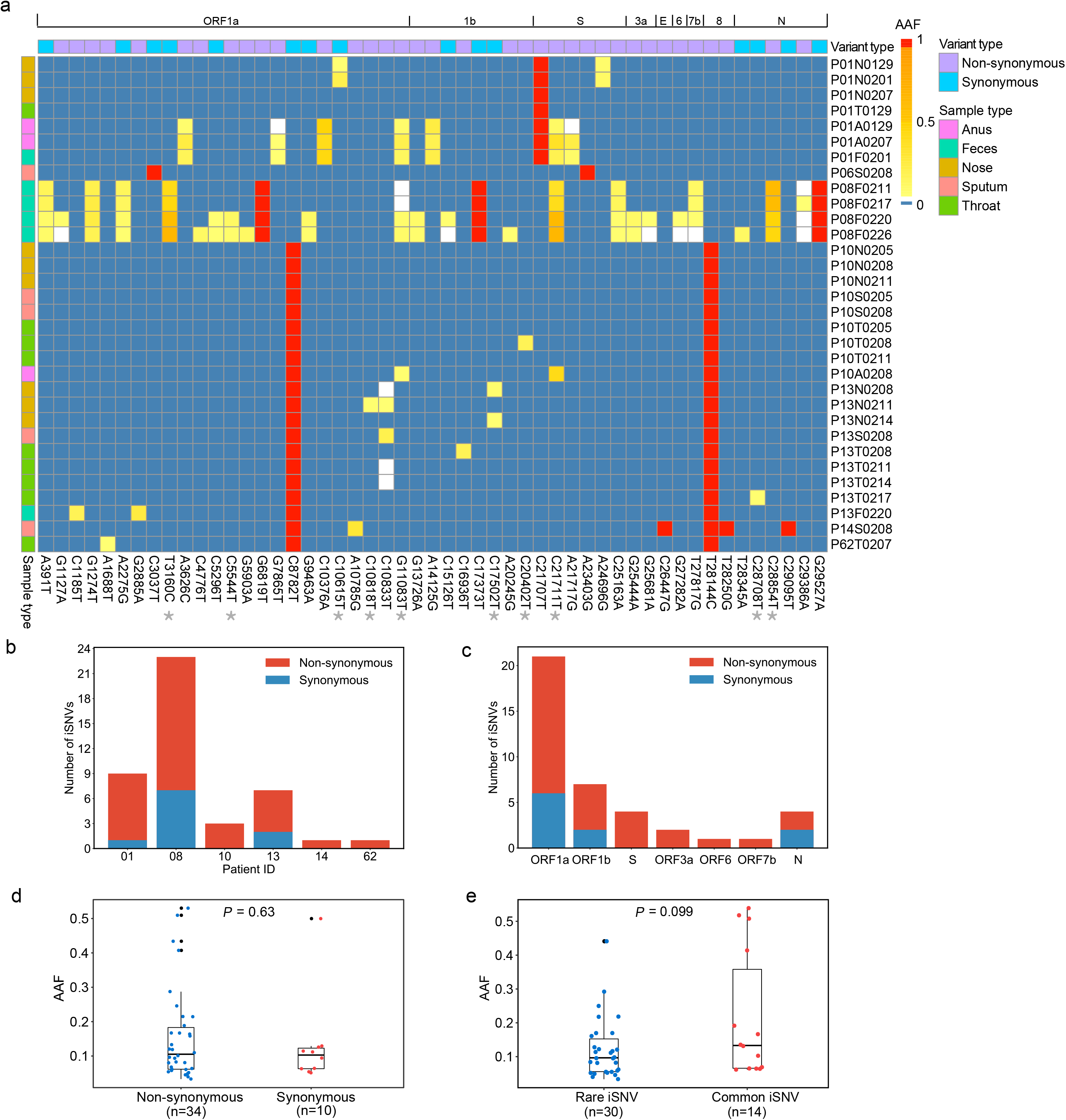
Characteristics of iSNVs. **a**, Heatmap showing the alternative allele frequencies (AAFs) of intra-host single nucleotide variants (iSNVs) and consensus variants among samples. The sample (e.g P01N0129) name indicates patient number P01, sample type (N nosal swab, T throat swab, A anal swab, F feces, S sputum) and collection date (01-27). **b**, The number of detected iSNVs per patient. **c**, Number of iSNV sites among protein-encoding genes. **d**, Box plot showing the distribution of alternative allele frequencies (AAFs) of non-synonymous and synonymous iSNVs. Each dot indicates the median AAF among all the detected iSNVs of samples from same patient. **e**, Box plot showing the distribution of AAFs of common and rare iSNVs. Each dot indicates the median AAF among all the detected iSNVs of samples from same patient.

We further analyzed intra-host variation across genes for evidence of natural selection. Overall, the 40 identified iSNV sites (10 synonymous iSNVs and 30 non-synonymous iSNVs) distributed evenly across genomic regions (**Fig. 2c; Table S3**). High proportion of non-synonymous iSNVs suggests that most iSNVs were either under frequent positive selection or insufficient purifying selection. However, we did not observe significant difference in AAFs between non-synonymous and synonymous iSNVs (**Fig. 2d**) and among codon positions (**Fig. S4**), indicating a relaxed intra-host selection. It is likely that most of those non-synonymous iSNVs will be removed by purifying selection and/or genetic drift in a longer timescale^6^. Nonetheless, the exact functional and evolutionary relevance of the intra-host variants remain to be explored.

One central task when estimating intra-host variation is to identify the source of iSNVs. Overall, the distribution of the iSNVs among samples does not correlate well with the consensus SNPs (**Fig. 2a**). Samples carrying the same consensus SNPs generally had different iSNVs, particularly in P01, P10 and P13. Here, we classified the iSNVs into i) rare iSNVs (30/40) detected in a single patient, and ii) common iSNVs (10/40) detected in at least two patients and/or identical to consensus variants. The ten common iSNVs did not show significant higher AAFs than the rare iSNVs (**Fig. 2e**). Notably, the ten common iSNVs include two iSNVs (G11083T and C21711T) exclusively detected in the GIT populations of P01, P08 and P10 (**Table S4**). Among the common iSNVs, G11083T is the most widespread consensus variant distributed in multiple lineages of SARS-CoV-2, suggesting that it might derive from recurring mutations on distinct strains rather than the mutation on a single ancestral strain. Interestingly, although G11083T was detected as an intra-host variant in the GIT samples of three patients, it was not detected in the corresponding RT samples, indicating a recurrent mutation of this loci, especially in the GIT population. Interestingly, G11083T locate in a region encoding a predicted T-cell epitope^7^, suggesting that recurrent mutation may provide genetic plasticity to better adapt against host defenses.

Using Shannon entropy, we observed a significantly higher genetic diversity within the GIT samples than that of RT samples (Wilcoxon rank-sum test, *P* = 1.4e-05; **Fig. 3a and Table S5**), reflecting an increased viral population size within the GIT samples. We further investigated the genetic differentiation between the two places. Notably, no iSNVs was shared between RT and GIT samples from the same patients, suggesting a clear genetic divergence among intra-host viral populations. Here we used L1-norm distance to estimate genetic dissimilarity among samples based on iSNVs and their AAFs and compared that between samples within and among hosts (**Fig. 3b and Table S6**). As expected, genetic distances among samples from the same host were smaller than those among inter-host samples (**Fig. 3b and Table S6**). Within each host, the greatest genetic differentiation was observed among GIT samples and between GIT and RT samples, while the differentiation among RT samples was relatively small. For example, seven iSNVs were shared among the GIT samples of P01, while none of them was observed in RT samples (**Fig. 2a**). It seems that the frequent genetic divergence between GIT and RT populations is mostly driven by bottleneck events during distant intra-host transmissions. However, the exact interaction mechanisms among intra-host populations require further investigation.

**Figure 3.**
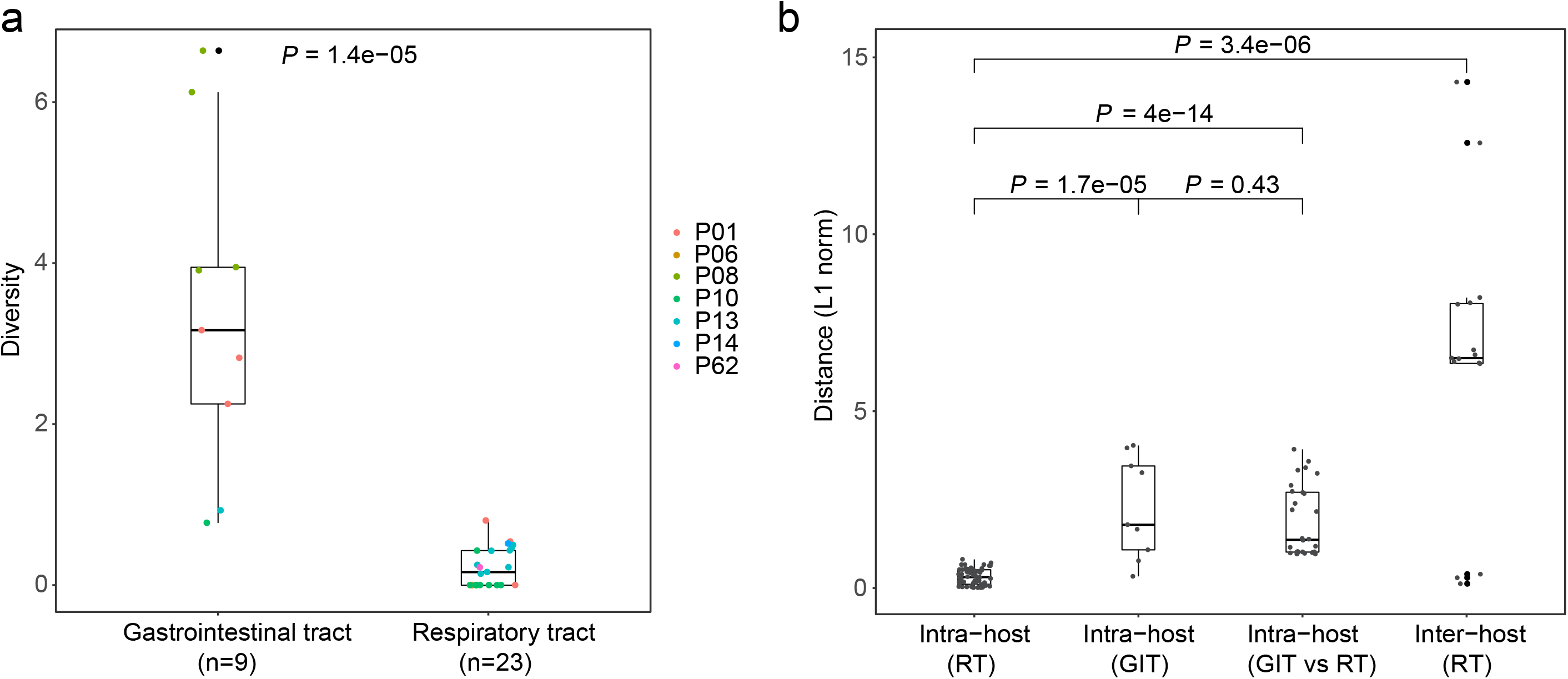
Dynamics of iSNVs detected in SARS-CoV-2 infected patients. **a**, Box plot showing the distribution of genetic diversity among samples from gastrointestinal tract (GIT) and respiratory tract (RT). **b**, Box plot showing the distribution of L1-norm distances among samples from gastrointestinal tract (GIT) and respiratory tract (RT). Each dot represents the genetic distance between a unique pair.

Previous studies have revealed longitudinal evolution of intra-host populations in some important RNA viruses^8–10^. We firstly compared the detected iSNVs among serial samples. All the iSNVs of early GIT samples also presented in later GIT samples, while all the iSNVs detected in RT samples disappeared in the following samples, suggesting that the viral genetic diversity is better maintained in GIT. We further focused on the allele frequency dynamics of GIT iSNVs of P01 and P08, respectively. Notably, most GIT iSNVs were remarkably stable and showed continuous trends of AAFs across sampling dates. For example, within the GIT population of P01, seven iSNVs showed continuous trends of allele frequency dynamics, including four iSNVs with increased AAFs and two iSNVs with decreased AAFs across the three sampling dates (**Fig. 4a**). Given their similar growth rates but distinct allele frequencies, it is likely that more than two genetically related haplotypes co-existed in within P01. Similar patterns were also observed in the GIT population of P08 (**Fig. 4b**). Notably, the dynamics of intra-host variants changed the consensus allele (>50%) of three genomic loci (3160, 21711 and 28854) of P08. Taken together, the iSNVs and their frequencies suggest that the viral populations in GIT is more stable than those in RT. Nonetheless, in both P01 and P08, we observed increased AAFs of C21711T and G11083T, suggesting that these two variants might be adaptively selected, especially in the GIT. Whether viral adaptation is involved in the intra-host divergence among distant populations warrants further investigation.

**Figure 4.**
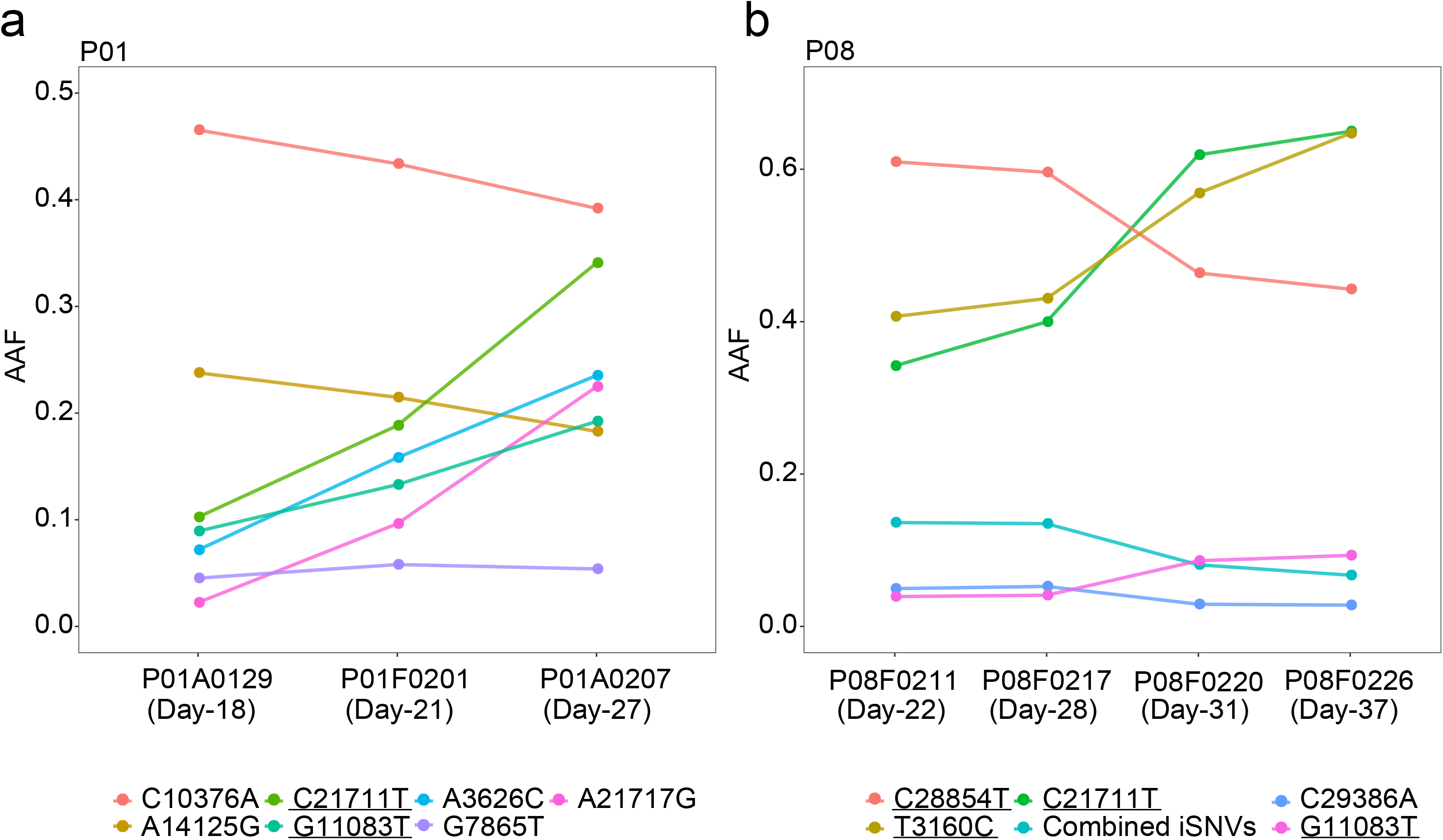
Temporal dynamics of intra-host populations in patient P01 and P08. **a-b**, Alternative allele frequencies (AAFs) among sampling dates in patient P01 and P08. Days post the first symptom date are shown in bracket. Combined iSNVs are the average frequency of four similar iSNVs (A391T, A2275G, C25163A and T27817G). Colours represent different iSNVs. Underlines represent common iSNVs.

We further phased the proximal iSNVs into local haplotypes using paired-end mapped reads (**Table S7**). Most minor haplotypes had one nucleotide difference from the dominant haplotype of the same sample, suggesting that they might derive from the main strain of the population. Nonetheless, we observed one exception in the GIT population of P01, covering the variable sites of C21707T, C21711T and A21717G (**Fig. S5**). With the cut-off of 1%, one dominant haplotype (T-C-A) and two minor haplotypes (T-T-A and T-T-G) were identified. Despite that minor haplotype (T-T-A) was relatively stable (8%–10%), the proportion of the dominant haplotype (T-C-A) decreased from 89% to 67%, while that of the minor haplotype (T-T-G) increased from 2% to 22%. Based on the dynamics and nucleotide differences among three haplotypes, we hypothesized that the minor haplotype (T-T-G) may derive from the dominant haplotype (T-C-A) via the intermediate haplotype (T-T-A), showing a maintained diversity within GIT population. More importantly, our observation supports that the mutated viruses are capable to replicate and hence, accumulate more variants within GIT of the same host, leading to an increased genetic diversity in the tissue specific environment.

Given the observations in patients with influenza^8^, stochastic process is the dominant factor driving the intra-host population dynamics, which is especially the case during distant intra-host transmissions. For SARS-CoV-2, one possible intra-host transmission route is from the respiratory tract to the gastrointestinal epithelia. During the intra-host transmission, population composition may change dramatically through random sampling when a novel sub-population was established from a small group of viruses of a larger population^13^. This is supported by the genetic divergence of intra-host variants between RT and GIT populations. The stochastic process between and within intra-host populations seems to also attenuate the efficacy of intra-host purifying selection, as shown by the even distribution of AAFs among synonymous and non-synonymous iSNVs. However, under the traditional genetic population theories, novel founder populations are expected to have a low genetic variation due to the subsampling from the original population. In contrast, viral populations in GIT showed a higher genetic diversity than those in RT, reflecting a larger effective viral population size in the GIT. This result is also consistent with the high viral load in GIT (**Fig. 1b**). During the viral replication, both RT and GIT populations showed evidence of generating intra-host variants. Our findings further demonstrated that those novel and/or recurrent intra-host variants are better maintained within GIT, and hence, leading to a higher level of genetic diversity and potentially larger effective population size in GIT. In contrast, the intra-host variants seemed to be less stable in RT, probably associated with a more dramatic genetic drift in RT populations. Differences in other factors, such as host-cell entry, immune responses and microbial communities among tissue specific environments, may further drive the structuring among intra-host population. On the other hand, those differences may also drive viral adaptation, given the two GIT specific non-synonymous iSNVs observed in our study. However, it is still challenging to fully disentangle the influences of stochastic processes and natural selection, considering the frequent confounding genetic signals of these two processes.

Intra-host variants were identified in many RNA viruses^8,9,11–14^. Here, using deep sequencing data of serial samples, we revealed the existence of intra-host variation within COVID-19 patients, which is likely to be contributed by novel and/or recurring intra-host mutations. Furthermore, our observation demonstrated a frequent genetic divergence between GIT and RT samples, mostly driven by bottleneck events among intra-host transmissions. Nonetheless, we observed a maintained viral genetic diversity within GIT, reflecting an increased population with accumulated mutations developed in the tissue-specific environments. Exact biological mechanisms of the intra-host population dynamics remain to be explored in future. Our data presented here also reflects the evolutionary capacity of SARS-CoV-2 in developing viral escape and drug resistance during infection. More broadly, these data provide new insights into the complex virus-host interactions.

## METHODS

### Patient enrollment and Ethics statement

Eight pneumonia patients, referred as GZMU cohort, were confirmed with SARS-CoV-2 infection between January 25 to February 10 in 2020 and hospitalized at the first affiliated hospital of Guangzhou Medical University (six patients), the fifth affiliated hospital of Sun Yat-sen University (one patient), and Yangjiang People’s Hospital (one patient). Serial samples were collected, including nasal swabs, throat swabs, sputum, gastric mucosa, urine, plasma, anal swabs and feces. The overall research plan was reviewed and approved by the Ethics Committees of all the three hospitals. All the information regarding patients has been anonymized.

### Real-time RT-qPCR and Metatranscriptomic sequencing

A total of 62 serial clinical samples collected from eight patients with COVID-19 (**Table S1**) were used for Real-time RT-qPCR. Clinical samples were subjected to RNA extraction using QIAamp Viral RNA Mini Kit (Qiagen, Hilden, Germany). An in-house real-time RT-qPCR was performed by targeting the SARS-CoV-2 RdRp and N gene regions (Zybio Inc.). Human DNA was removed using DNase I and RNA concentration was measured using Qubit RNA HS Assay Kit (Thermo Fisher Scientific, Waltham, MA, USA). DNA-depleted and purified RNA was used to construct double-stranded DNA library using MGIEasy RNA Library preparation reagent set (MGI, Shenzhen, China) following the protocol described in our previous study^15^. High throughput sequencing of the constructed libraries was then carried out on the DNBSEQ-T7 platform (MGI, Shenzhen, China) to generate metatranscriptomic data of 100bp long paired-end reads.

### Hybrid capture-based enrichment and sequencing

For a subset of samples (**Table S1**), genomic content of SARS-CoV-2 was enriched from the double-stranded DNA libraries mentioned above using the 2019-nCoVirus DNA/RNA Capture Panel (BOKE, Jiangsu, China) as described in our previous study^15^. The SARS-CoV-2 content enriched samples were used to construct DNA Nanoballs (DNBs) based libraries, which were then sequenced using the same protocol described above.

### Data filtering and Genome assembly

Data filtering was performed following the procedures described in previous research^15^. Briefly, for both metatranscriptomic and hybrid capture data, sequence data of each sample were firstly mapped to a pre-defined database comprising representative genomes of coronaviridae. The mapped reads were then subject to the removal of low-quality, duplications, adaptor contaminations and low-complexity to collect high quality coronaviridae-like reads. We also compared the allele frequencies among the two data types when available, samples with conflicted consensus alleles were removed. For the samples with 60-fold of metatranscriptomic data, coronaviridae-like metatranscriptomic reads were used to generate consensus genomes and identify intra-host variants. Full-length consensus genomes were generated from reads mapped to the reference genome (GISAID accession: EPI_ISL_402119) using Pilon (v. 1.23)^16^. To prevent false discovery, base positions reporting an alternative allele with the following conditions were masked as N: 1) sequencing coverage less than 5-fold; 2) sequencing coverage less than 10-fold and the proportion of reads with the alternative allele less than 80%. The collected coronaviridae-like reads were also de novo assembled using SPAdes (v. 3.14.0) with default settings^17^ with a maximum of 100-fold coverage of read data. Structural variations between the de novo assemblies and consensus genomes, if any, were manually checked and resolved based on read alignments. Nucleotide differences between the consensus sequences and the reference genome were summarized into artificial Variant Call Format (VCF) files, which were annotated using SnpEff (v.2.0.5)^18^ with default settings.

### Phylogenetic analysis

Available consensus sequences of SARS-CoV-2 (**Table S8**) were collected from GISAID database (https://www.gisaid.org/) on 5th April, 2020, after the removal of highly homologous sequences, 122 representative virus strains (**Table S8**) were used to infer evolutionary relationships with the assembled genomes. Within the GZMU cohort, only one genome was selected when more than one identical genome was achieved from the same patient. The assembled SARS-CoV-2 and selected representative genomes were aligned using MAFFT with default settings. A maximum likelihood (ML) tree was inferred using the software IQ-TREE (v.1.6.12)^19^, with the best fit nucleotide substitution model selected by ModelFinder from the same software. The inferred ML tree was then visualized using the R package ggtree^20^ (v.3.10). Major branches and the defining nucleotide mutations were manually labelled.

### Summary of public consensus variants

All the consensus sequences of the public strains were aligned with the reference genome (GISAID accession: EPI_ISL_402119) using MAFFT (v.5.3)^21^ with default settings. Nucleotide differences between the consensus sequences and the reference genome were summarized into an artificial VCF file, which was then were annotated using SnpEff (v.2.0.5) with default settings. The linkage disequilibrium among the identified consensus variants were estimated using VCFtools (v.0.1.16).

### Calling of iSNVs

Here, an intra-host single nucleotide variant (iSNV) was defined as the alternative allele co-existed with the reference allele at identical genomic position within the same sample. To minimize false discovery, iSNVs were identified on samples with at least 60-fold mean metatranscriptomic sequencing coverage and then verified using hybrid-capture data when available.

First, paired-end metatranscriptomic reads were mapped to the reference genome (GISAID accession: EPI_ISL_402119) using BWA aln (v.0.7.16) with default parameters^22^. Duplicated reads were marked using Picard MarkDuplicates (v. 2.10.10) (http://broadinstitute.github.io/picard) with default settings. Base composition of each position was summarized from the mapped reads using the software pysamstats (v. 1.1.2) (https://github.com/alimanfoo/pysamstats), and then subject to iSNV site identification with following criteria: 1) base quality larger than 20; 2) sequencing coverage of paired-end mapped reads >= 10; 3) at least five reads support the minor allele 4) minor allele frequency >= 5%; 5) strand bias ratio of reads with the minor allele and reads with major allele less than ten-fold. To minimize false discoveries, sites with more than one alternative allele were filtered out. Biological effects of the identified iSNVs were annotated using the SnpEff (v.2.0.5) with default settings. Alternative allele frequencies (AAFs) at the identified iSNV sites were measured by the proportion of paired-end mapped reads with alternative alleles. When an iSNV was detected in one patient, the detection cut-off of that iSNV was reduced to 2% for the rest samples of the same patient. Only the AAFs more than 2% with at least three reads were kept for the following analyses. All the iSNVs were verified using hybrid capture data when available. At the iSNV sites, the allele with higher frequency was defined as major allele, while one with less frequency was defined as minor allele, regardless whether it is different from the reference allele. A heatmap was generated to visualize the AAFs for all samples using the pheatmap package in R (v.3.6.1). A subset of the identified iSNVs were validated by Sanger sequencing using the protocol described in previous study^15^.

### Statistics of iSNVs

The distribution of iSNVs among genetic components and patients were summarized and visualized using the Python package matplotlib (v.3.2.1). Alternative allele frequencies on all the detected iSNV sites were compared among patients. To avoid oversampling, for the patient with more than sample, only the median AAF among all samples of that patient was used for comparison. Alternative allele frequencies among synonymous and nonsynonymous variants and among codon positions were compared using Wilcoxon rank sum test and visualized through boxplot using the R package *ggplot* (v.3.3.0). For the iSNVs detected in patient P01 and P08, the dynamics of AAFs was visualized across time points using the R package *ggplot* (v.3.3.0).

### Genetic diversity

Genetic diversity of each sample was estimated using Shannon entropy based on the AAF of each iSNV, assuming that all iSNVs are independent from each other.

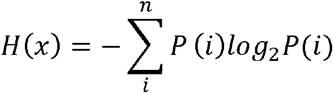

where *P*(*i*) is the AAF at variable site *i*.

### Genetic distance

The genetic distance among samples was estimated using L1-norm distance in a pairwise manner.

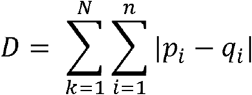

The L1-norm distance (*D*) between a pair of samples is the sum of distance across all the variable sites (*N*) For each variable site, the distance is calculated between vectors (*p* and *q* for each sample) comprising frequencies of all the four possible nucleotide bases (*n* = 4).

### Haplotype reconstruction

Haplotypes of neighbor iSNV sites were reconstructed using mapped paired end reads.

## DATA AVAILABILITY

Sequence data used in this study have been deposited in CNGB (https://db.cngb.org/) under Project accession CNP0001004 and CNP0000997.

## ACKNOWLEDGEMENTS

This study was approved by the Health Commission of Guangdong Province to use patients’ specimen for this study. This study was funded by grants from The National Key Research and Development Program of China (2018YFC1200100, 2018ZX10301403, 2018YFC1311900), the emergency grants for prevention and control of SARS-CoV-2 of Ministry of Science and Technology (2020YFC0841400) and Guangdong province (2020B111108001, 2018B020207013, 2020B111112003), the Guangdong Province Basic and Applied Basic Research Fund (2020A1515010911), Guangdong Science and Technology Foundation (2019B030316028), Guangdong Provincial Key Laboratory of Genome Read and Write (2017B030301011), Guangzhou Medical University High-level University Innovation Team Training Program (Guangzhou Medical University released [2017] No. 159), National Natural Science Foundation of China (81702047, 81772191, 91842106 and 8181101118), State Key Laboratory of Respiratory Disease (SKLRD-QN-201715, SKLRD-QN-201912 and SKLRD-Z-202007). We thank the authors for submitting the genome sequences to GISAID. We thank the Guangdong Provincial Key Laboratory of Genome Read and Write and China National GeneBank at Shenzhen for providing sequencing service. We thank the patients who took part in this study.

## AUTHOR CONTRIBUTIONS

J.Z., J.L., Y.L and J.Z conceived the study, Y.W et al collected clinical specimen and executed the experiments. D.W., W.S., X.C. and J.J. analyzed the data. All the authors participated in discussion and result interpretation. D.W., Y.W., M.P. and J.Z. wrote the manuscript. All authors revised and approved the final version.

## SUPPLEMENTARY INFORMATION

**Figure S1. Correlation of estimated alternative allele frequencies between biological replicates.**

**Figure S2. Correlation of estimated alternative allele frequencies between metagenomic and hybrid capture data.**

**Figure S3. Correlation between sequencing depth and detected iSNVs.**

**Figure S4. Number of iSNV among three codon positions.**

**Figure S5. Haplotype frequency of proximal iSNVs within the gastrointestinal tract of the patient P01**

**Table S1. Summary of clinical samples and patients with COVID-19**

**Table S2. Genomic information of 32 SARS-CoV-2 samples**

**Table S3. List of intra-host single nucleotide variants within 32 SARS-CoV-2 samples**

**Table S4. Allele frequency of iSNVs detected from metatranscriptomic and/or hybrid capture data**

**Table S5. Genetic diversity of 32 SARS-CoV-2 sampes**

**Table S6. Genetic distance between paired samples**

**Table S7. Frequency of proximal iSNVs using paired-end mapped reads**

**Table S8. List of public genomes used for analysis**

