## Supplementary figures and images for "Intra-host Variation and Evolutionary Dynamics of SARS-CoV-2 Population in COVID-19 Patients"

### Supplemental Figure S1

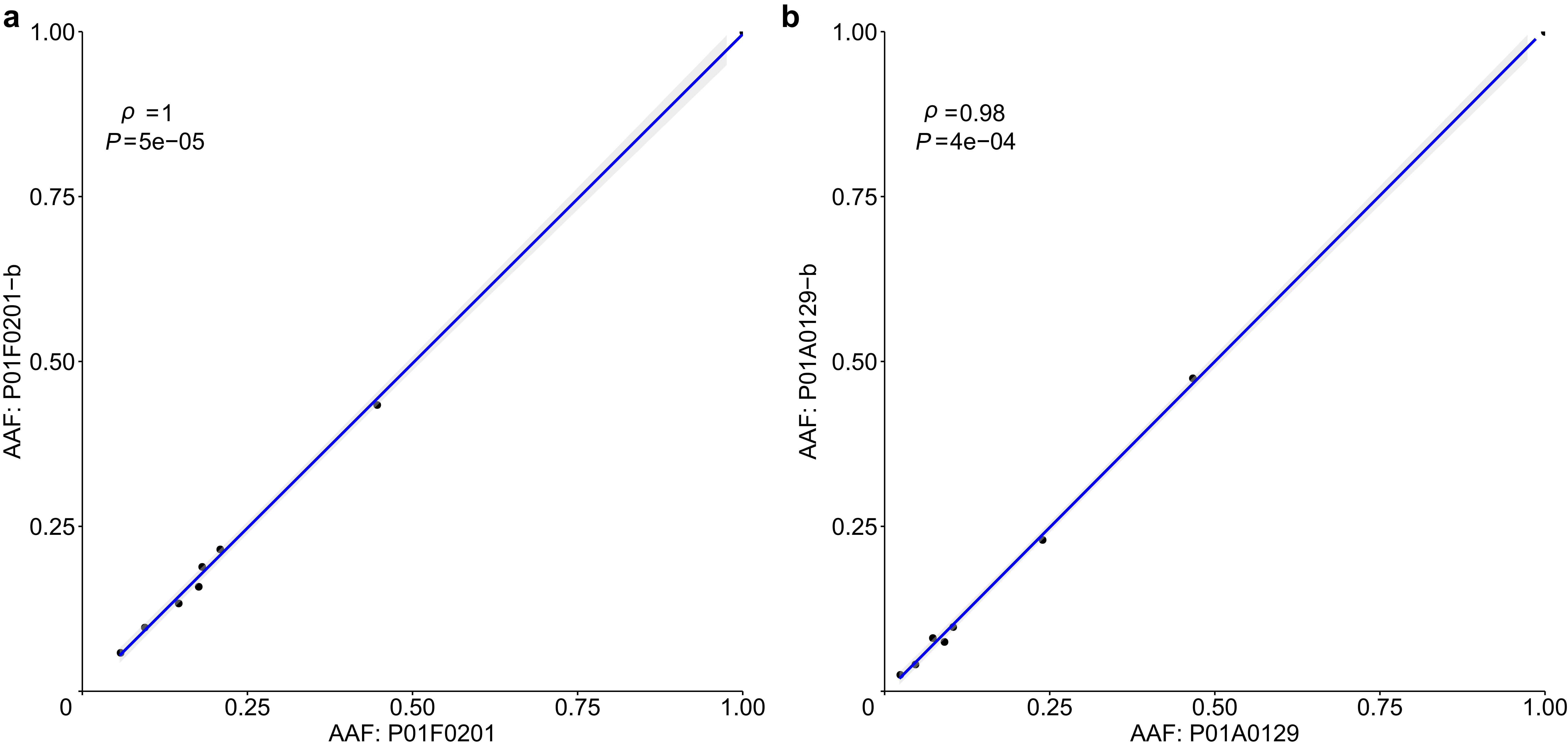

### Supplemental Figure S2

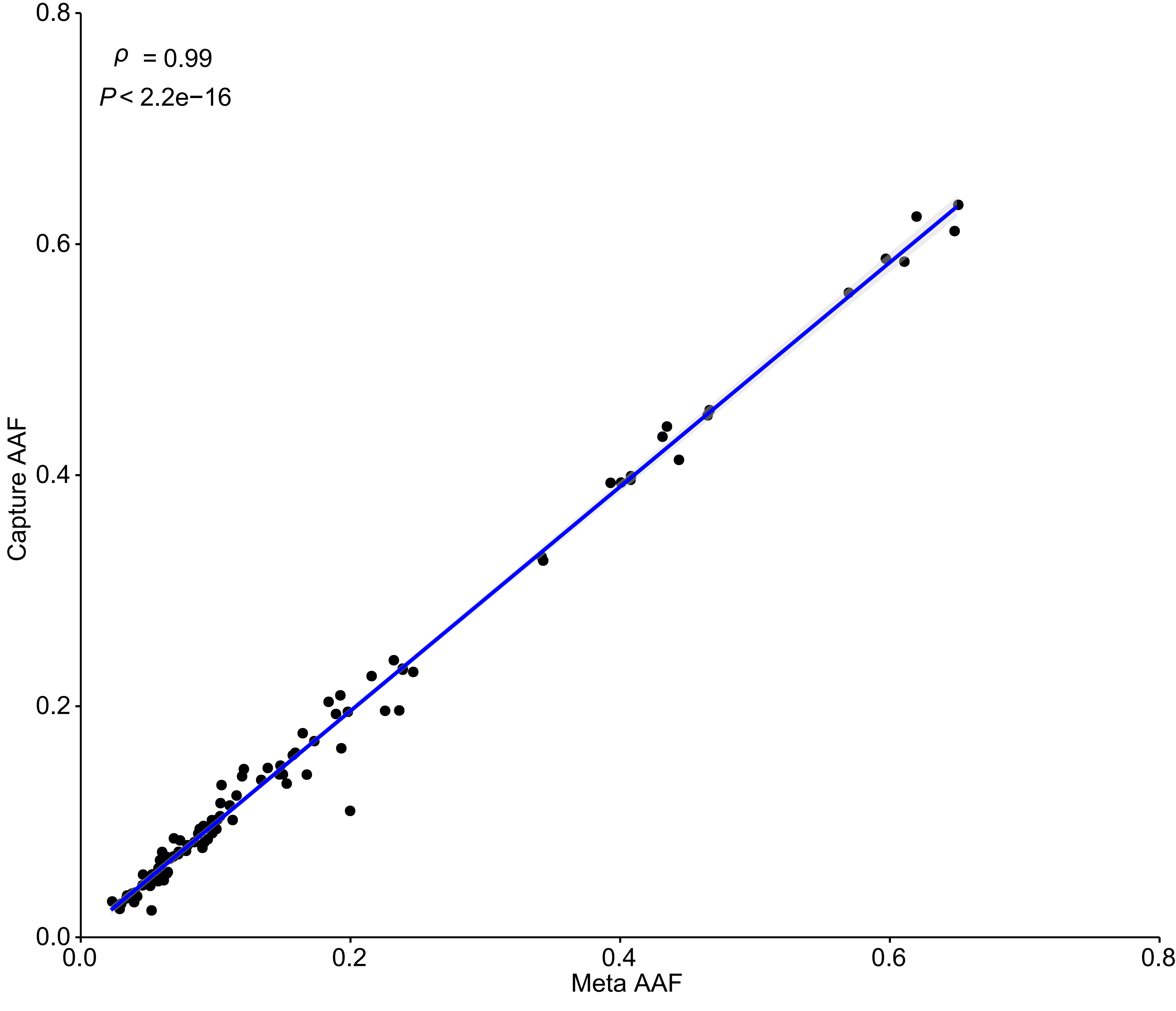

### Supplemental Figure S3

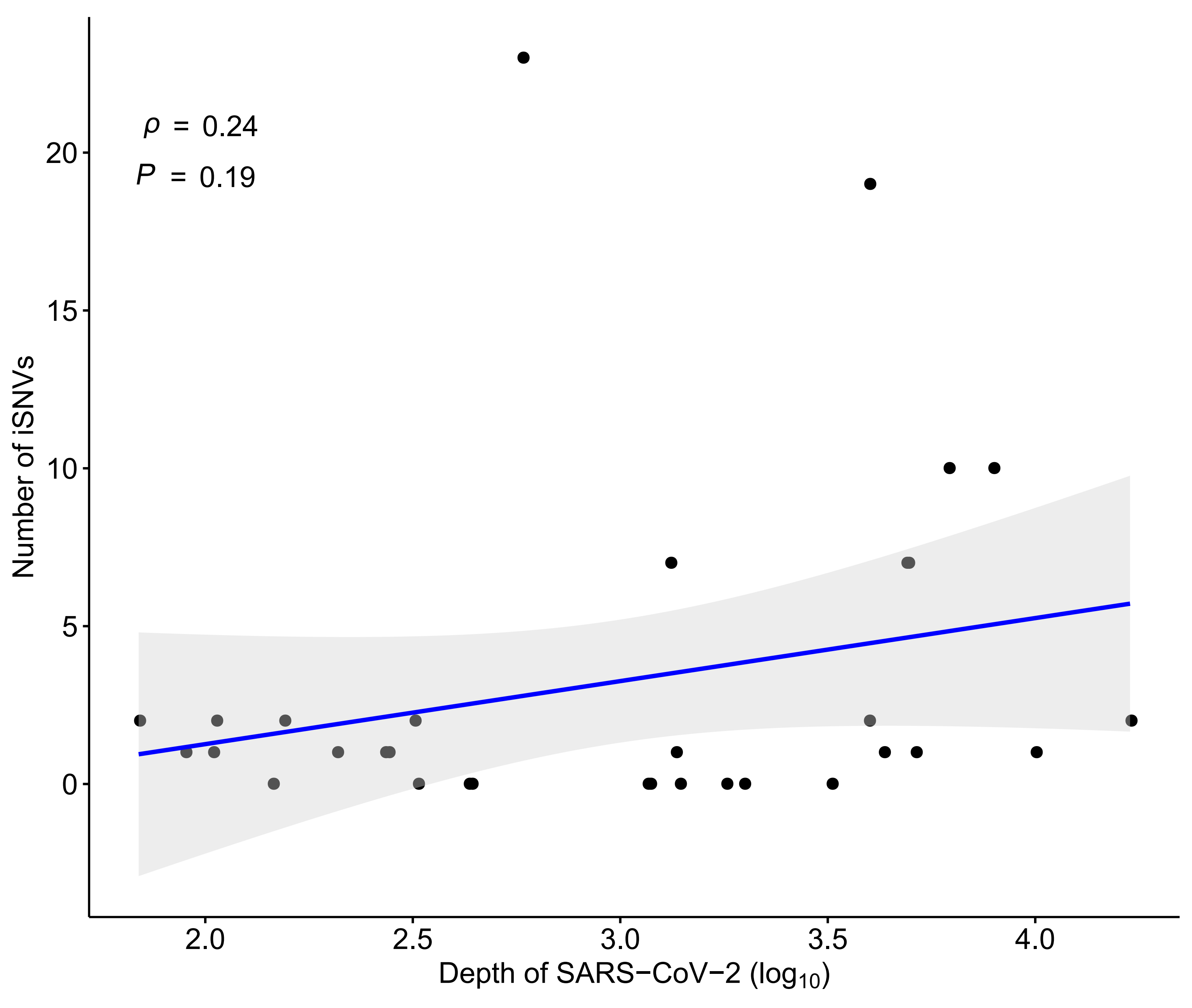

### Supplemental Figure S4

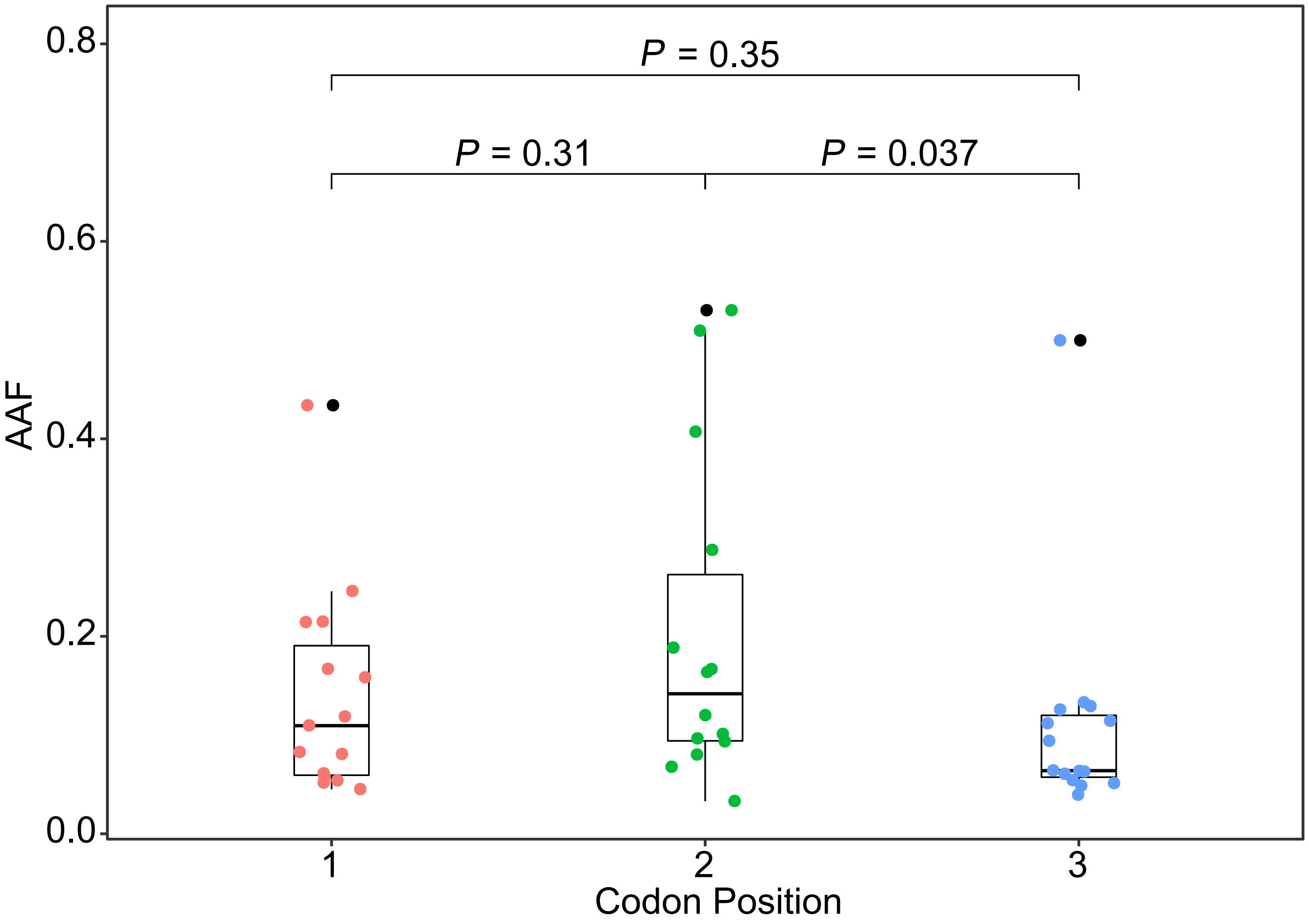

### Supplemental Figure S5

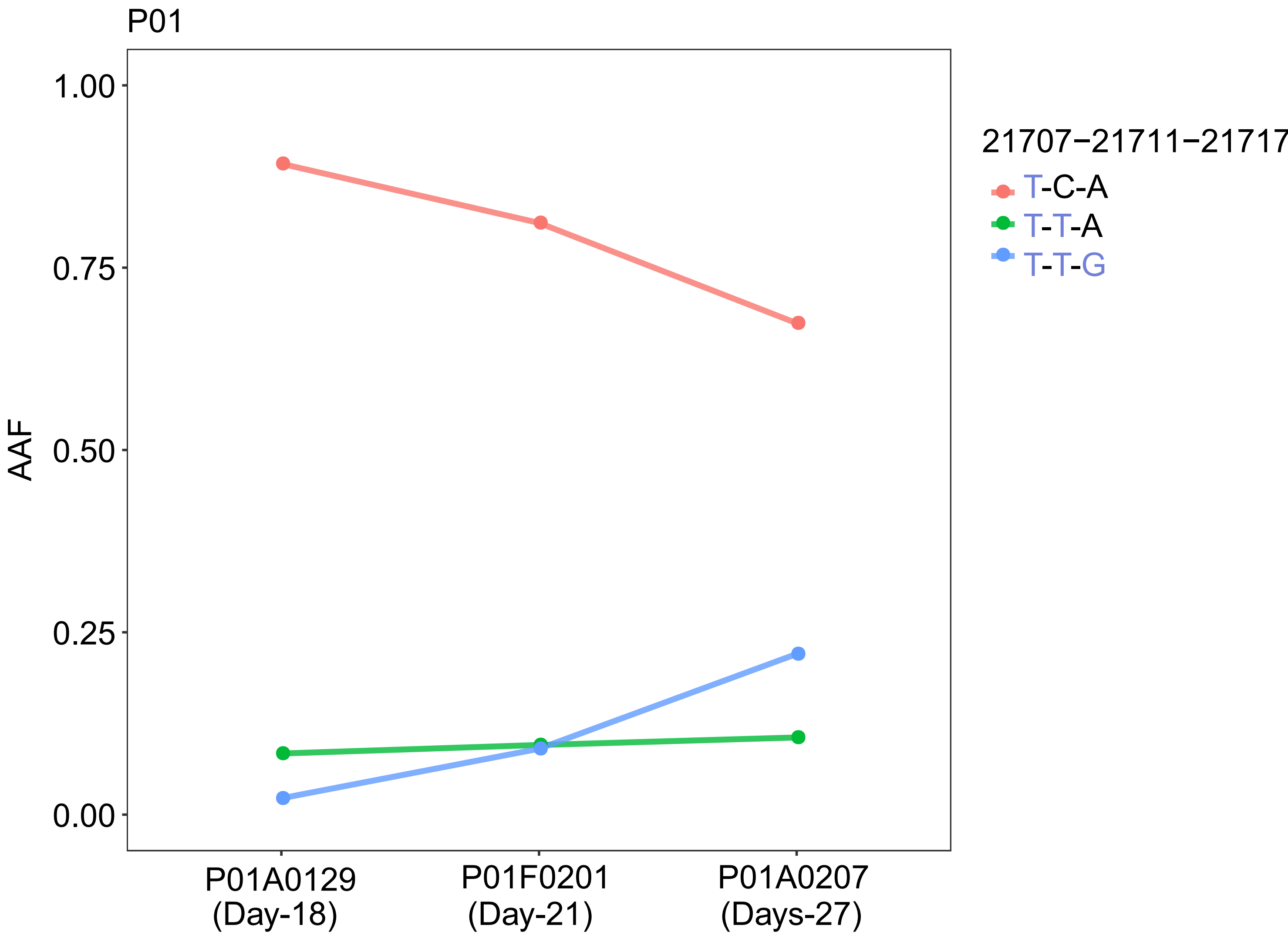
